# Sustaining yam yields amidst climate threat in the forest – savannah transition zone of Ghana

**DOI:** 10.1101/474247

**Authors:** F. Frimpong, Danquah E. Owusu, S. A. Ennin, H. Asumadu, A.K. Aidoo, N. Maroya

## Abstract

With about 70% of yam tuber been water, yield is critically affected during bulking as a result of onset of temporal drought. As a consequence of climate change, farmers who are into *Dioscorea rotundata* (white yam) production for local and international market lose their investments mainly due to erratic precipitation, drought spells culminating into low yields of just 12t/ha compared to the potential of about 22-49t/ha depending on the variety. Innovative land uses technologies with higher and sustained productivity for yam production are imperative. This study verifies improved agronomic package for sustainable yam production in yam growing areas in the forest – savannah transition zone of Ghana during the 2015 and 2016 cropping seasons. The improved agronomic package included use of ridging as seedbed, seed treatment before planting, fertilizer application at a rate of 30:30:36 N:P_2_0_5_:K_2_0 kg/ha plus 15 kg/ha Mg and 20 kg/ha S as MgSO_4_ and the use of minimum stakes (trellis; 30-50% less number of stakes used by farmers staking). This was compared with farmers’ practice which consisted of mounding, no fertilizer application and no seed treatment. The results revealed significant (*P* ≤ 0.01) yam yields of more than 60% difference between the improved agronomic practice and farmers’ practice from Ejura, Atebubu and Kintampo yam growing communities. Adoption of improved agronomic practices does not only sustain yam production and address deforestation but also provide higher returns on investments promoting climate resilience by small holders.

## Background

In Ghana, crops are already experiencing heat stress, drought spells, several pests and diseases outbreak and shorter growing crop duration as a consequent of the changing climate (1). Thus, a potential catastrophe for smallholder farmers and the millions of people who regularly grow rain-fed full season crops such as yam, cocoyam, rice etc.(2–5). Choices about what to grow are often dictated by the ability of the rainfall regime to support moisture for plant growth (6). One way around this would be to breed for varieties with shorter crop maturity durations or management interventions that build on the resilience of cropping production systems to reduce shocks if the shocks from the climate change cannot be done away with. Evidence suggests that climate smart agriculture can make a contribution to mitigation by supporting more efficient use of fertilizers, weed management and reduced staking options in yam production (1,7,8).

Yam, an important staple food crop across West Africa is a major non-traditional export crop in Ghana contributing to about 16% of the National Agricultural Gross Domestic Product (9,10). However, there are a number of challenges that hamper the production and productivity of yam. Predominantly amongst them are; inadequate rainfall, low soil fertility, weed infestation, pests and diseases in the field (foliar and soil borne) and inadequate storage facilities, attack by organisms such as rodents, access to quality improved seed, implements for mechanization etc.(8,11). Others include shortage of stakes especially from deforested areas and guinea savannah regions. This is as a consequent of clearing new lands year after year popularly known as shifting cultivation and slash and burn agriculture, inadequate labour in view of the labor intensive nature of yam cultivation (e.g. for preparation of mounds, staking, harvesting)(8,12,13). This current yam production system where there is annual shifting of farm to new lands is not sustainable and therefore the urgent need to disseminate an environmentally sound yam production technology that would increase yield and sustain production on continuously cropped fields particularly in the face of climate change (14).

As a follow up on an on-station evaluation conducted in 2013 and 2014 cropping seasons at the CSIR-Crops Research Institute research stations, recommended agronomic package of planting treated yam seeds on ridges with fertilizer rate of 30:30:36 N: P_2_0_5_:K_2_0 kg/ha plus 15 kg/ha Mg and 20 kg/ha S as MgSO_4_ and trellis staking were verified by comparing it with the farmers practices on farmers’ fields at Ejura, Atebubu and Kintampo yam growing communities. The use of ridges and yam seed treatment helps to maintain optimum number of stands per unit area and fertilizer addresses the soil nutrient depletion whiles the trellis staking option uses ropes and few stakes to address the challenge of scarcity of stakes and cutting of more trees/bamboo for staking. The study has a major objective of validating and demonstrating improved yam production technologies (macronutrients (NPK) and micronutrients (Mg & S), minimum staking option and seed treatment) to major yam growing communities in Ghana.

## Methodology

### Study sites characteristics

The experiments were conducted in 2015 and 2016 cropping seasons at Ejura-Sekyere dumasi (Aframso/Teacherkrom, Ashakoko, Dromankuma & Nkwanta), Atebubu-Amantin (Adom, Dagaati Line, Munchunso & Nwowamu) and Kintampo North (Asantekwa, Suamre, Babaso/Yabraso & Kintampo Magazine) districts of Ghana (Fig 1). These areas lies in the forest-savannah transition agro-ecological zone and amongst the major yam growing areas of Ghana (15). Eight farmers (4 randomly selected for analysis) from each of the three operational areas were selected for the study every season (Table 1). Mean annual rainfall (mm) for 2015 & 2016 across locations are shown in the map (Fig 1). The data were sourced from the local district weather stations which revealed a reduced rainfall amount (mm) in 2016 compared to 2015 with Kintampo communities severely affected (Fig 1). Mean annual rainfall (mm) pairs recorded for 2015 and 2016 were 1256:1034, 929:769, and 863:795 at Ejura, Atebubu and Kintampo respectively (Fig 1). These locations have bimodal rainfall i.e. major rainy season from March-Mid August and minor rainy season from September-November; peak in October. Temperature ranges from 25-39 C with soil type of Ferric Acrisol; Asuansi series, upper top soil consist of 5cm grayish brown sandy loam topsoil of dark brown gritty clay loam (16).

**Table 1:**
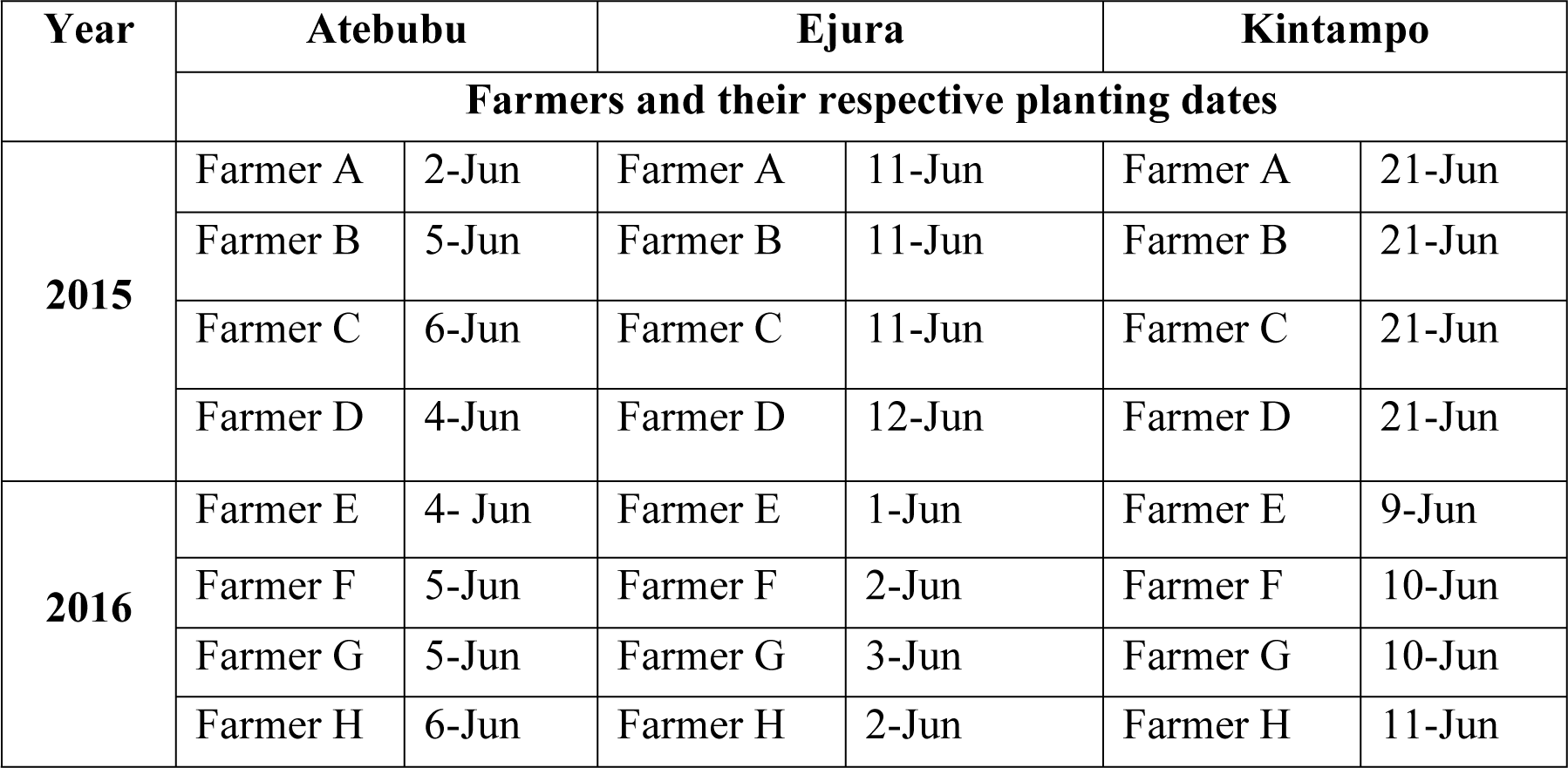
Farmers and planting dates selected from each of the operational areas for the analysis in 2015 and 2016.

**Figure 1.**
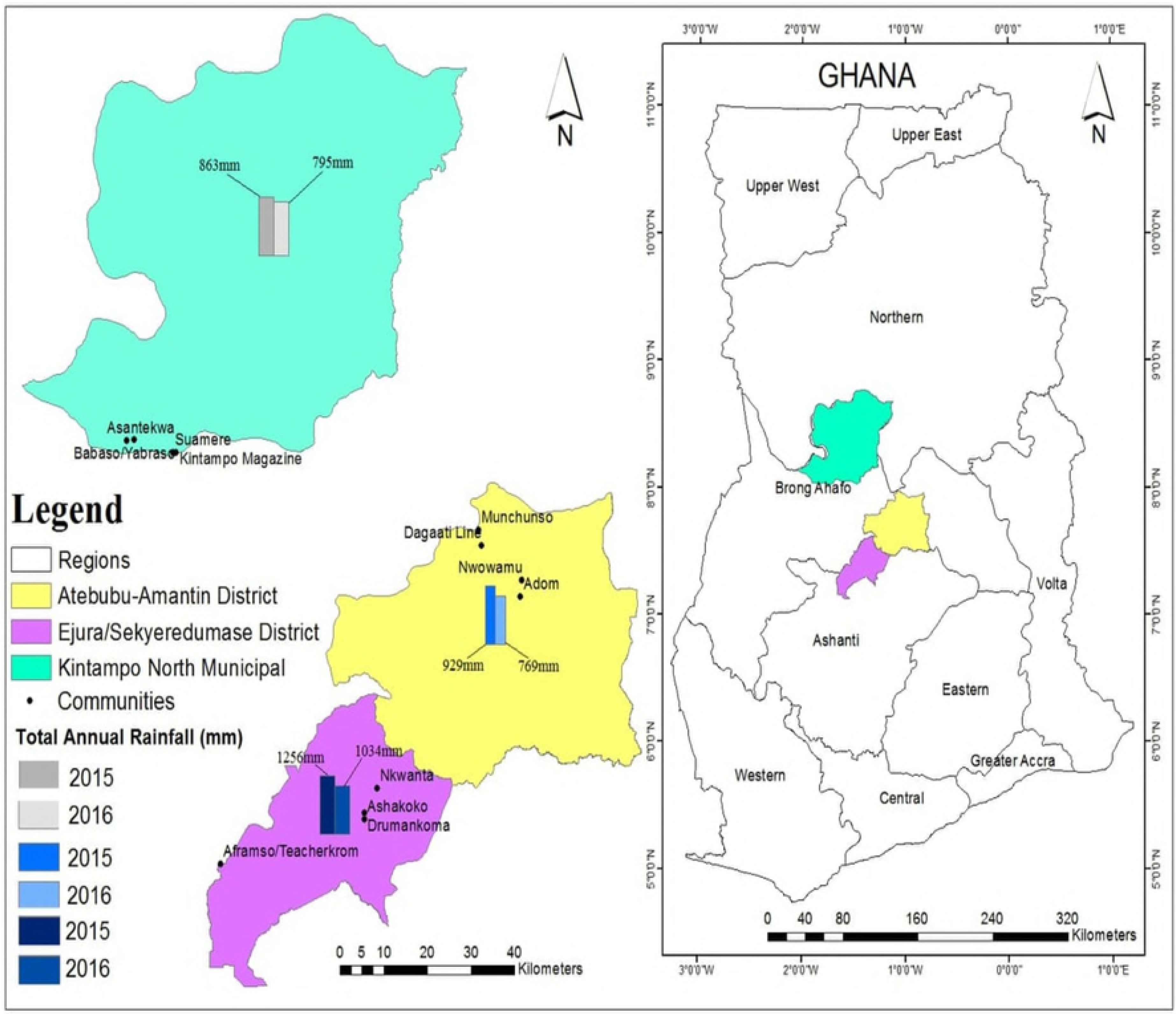
Map of Ghana on the (right), zoomed in on Ejura-Sekyeredumasi, Atebubu - Amantin and Kintampo North districts (left) of the forest-savanna transition zone. Farming communities where the studies were undertaken for 2015 and 2016 cropping seasons are illustrated with dots (.) with their names beside. Mean annual rainfall (mm) per location are depicted by bar plots for each season.

### Experimental design

A randomized complete block design with each farmer as a replicate was used for the study. A total of 8 trials (8 replicates) were established in all the operational areas in 2015 and 2016. Local white yam variety "*Dente"* of *Dioscorea rotundata* species was planted and subjected to two treatment applications from start of planting till harvest. Planting of yam across these locations were started in June and completed by 21-June of each year (Table 1). Harvesting of yam was completed by the end of December of each year. The treatments were recommended (improved agronomic practice) and local technology/farmers’ practice. The recommended practice for yam production included a package of; treating yam seed before planting with fungicide and insecticide, use of ridging as seedbed, fertilization at 30:30:36 N: P_2_0_5_:K_2_0 Kg/ha plus 15Kg/ha of Mg and 20Kg/ha S as MgSO_4_ and use of trellis for staking whiles in the farmers’ practice, farmers were allowed to use their local technology (planting on mounds), no pre-treatment of seed, no fertilizer application for comparison. Continuously cropped fields that farmers would normally not use for yam production were selected in each operational area for the study. Each improved agronomic field had an area of a quarter of an acre (0.25ac/0.1ha) planted at 1.2m inter row and 0.8m on the ridges (10,416 stands/ha).

The same size (0.25ac/0.1ha) was demarcated for the farmers practice treatment where they mounded sparsely to cover the entire field (3,400 - 4,000 stands/ha). Each farmer field was considered as a replicate and analysis compiled for each district/operational area together. The Fertilizer treatment was applied at 50% split at 5-6 weeks and 11-12 weeks after planting in all the locations. The seed setts (200-250g) of the improved agronomic fields were treated with Dursban (Chlorpyrifos from Dow Agro Sciences; 1.25 l/ha) and Mancozeb (Dithiocarbamate from Ag-Chem Africa 80%; 75 g in 15 l of water) before planting. Farmers’ sett sizes used ranged between 350g to 650g depending on each farmer. Emerged weeds in the improved agronomic fields were controlled with glyphosate (2.5 liter per ha) before the sprouting of the yam while farmers only slashed on their fields. There after weeds were manually controlled with cutlass and hoe in either improved agronomic field or farmers’ field by hoeing. In 2016, sensory evaluations were conducted with 50 participants (who were mainly farmers from the localities) in all areas after eating boiled yam during the December harvest. Fertilized and unfertilized boiled yam (coded at the blind side of the participants) using one on one questionnaire interviews, farmers scored for taste, texture, aroma and acceptability (Table 3).

**Table 2.**
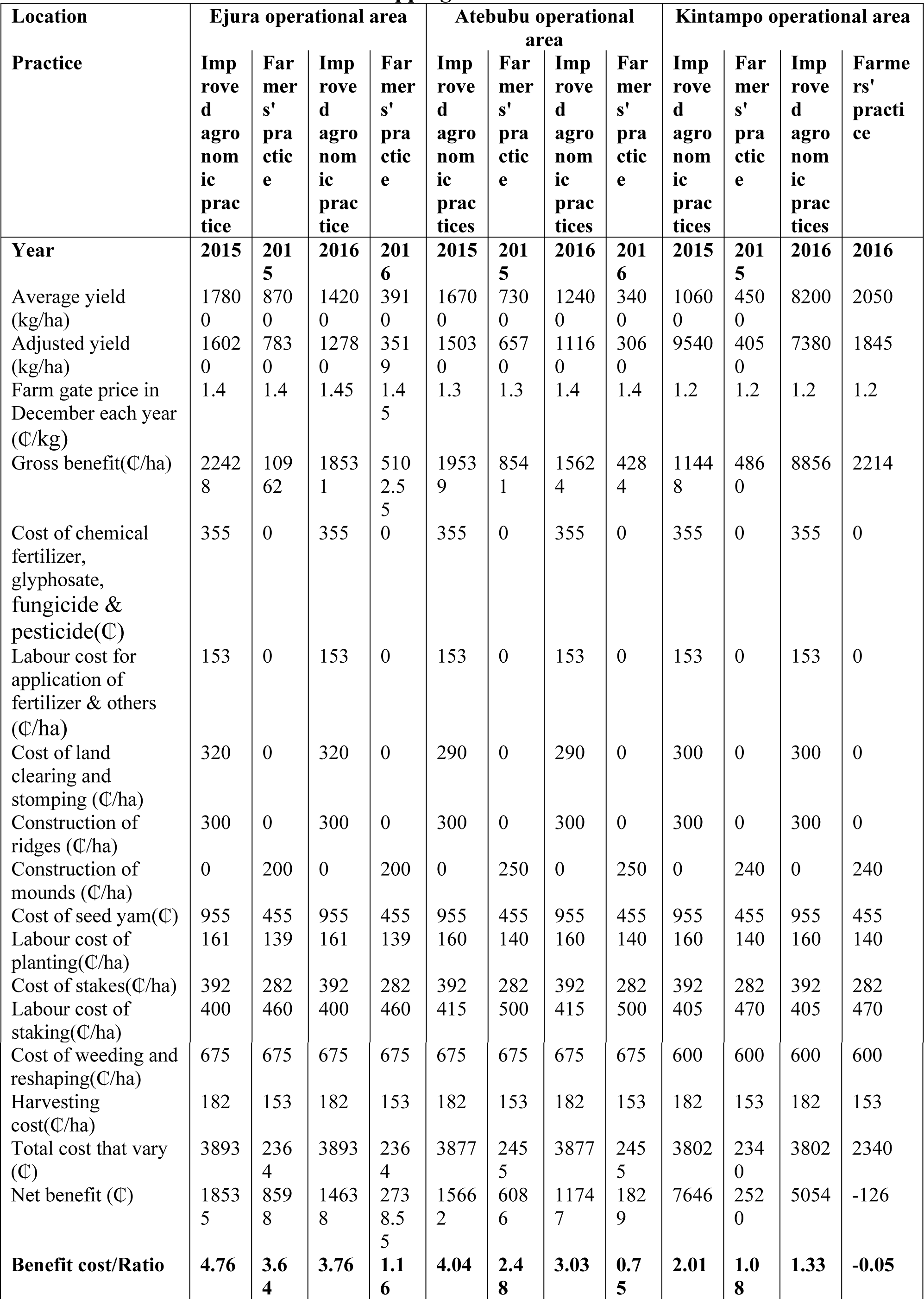
Partial budget and cost benefit analysis of white yam production with improved technology and farmers practices at Ejura, Atebubu and Kintampo farming communities for the 2015 and 2016 cropping seasons.

**Table 3:** Sensory evaluation of fertilized and unfertilized boiled yam involving farmers from Atebubu, Ejura and Kintampo farming communities.

| Location | Treatment | Taste | Texture | Aroma | Overall acceptability | STD acceptability |
| --- | --- | --- | --- | --- | --- | --- |
| Atebubu | Fertilized yam (30 30 36 N-P<sub>2</sub>O<sub>5</sub>-K<sub>2</sub>O (Kg/ha) + 20kg S & 15kg 15 Mg as MgSO<sub>4</sub>) | 2.30 | 2.5 | 2 | 2.20 | 0.87 |
|  | Unfertilized yam (0kg/ha) | 2.60 | 2.5 | 2.4 | 2.50 | 0.86 |
| Ejura | Fertilized yam (30 30 36 N-P<sub>2</sub>O<sub>5</sub>-K<sub>2</sub>O (Kg/ha) + 20kg S & 15kg 15 Mg as MgSO<sub>4</sub>) | 2.90 | 2.9 | 2.8 | 2.80 | 0.91 |
|  | Unfertilized yam (0kg/ha) | 2.90 | 2.9 | 2.8 | 2.80 | 0.91 |
| Kintampo | Fertilized yam (30 30 36 N-P<sub>2</sub>O<sub>5</sub>-K<sub>2</sub>O (Kg/ha) + 20kg S & 15kg 15 Mg as MgSO<sub>4</sub>) | 2.10 | 2.2 | 2.4 | 2.20 | 0.81 |
|  | Unfertilized yam (0kg/ha) | 2.70 | 3.3 | 3.1 | 3.10 | 0.97 |

*n=50, Score Scale 1-5; 1=best, 5=worst*

### Data collection and analysis

During harvest, the tubers were grouped into two; ware yam (tuber sizes of more than 500g) and seed yam (500g and below) and weighed separately for each of the practices. Four replications from each operational area (Ejura, Atebubu and Kintampo) and across the two seasons (2015, 2016) were subjected to statistical analysis. Data on stand harvested, weight of ware yam, weight of seed yam and total yam yield collected were subjected to one way analysis of variance linear model at 5% significant level using ‘R’ statistical software with the practice treatment as the independent variable. Where treatment means differ, Tukey’s HSD test was used to group them and visualized with bar graphs using MS excel 2010. Percentage differences of total yam yield harvested between the improved practice and the farmer’s was calculated based on the formula;

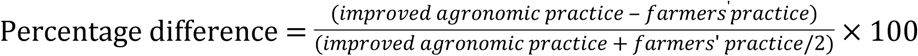

Sensory evaluation through one on one interviews upon tasting boiled yam from fertilized and unfertilized samples were calculated with data from 50 participants for each location who were mainly farmers. In order to deduce return on investment after venturing in any of the practices was subsequently calculated using benefit cost ratio for each of the locations using total yam yield and price per kg of yam at the time of harvest.

## Results and Discussion

### Influence of improved agronomic technology on yam tuber yield

Generally, total yam tuber yields were high in 2015 than in 2016 cropping season irrespective of the practice (Figs 2–4). The improved agronomic practice fields had significantly (P<0.01) higher total tuber yields compared to the farmers’ practice fields across the locations despite the season (Figs 2–4). Availability of moisture is critical for yam to sprout and establish during early growth stages and vital to bulk bigger tubers(11,17–20). Higher rainfall during 2015 cropping season (Fig 1) might have ensured yam establishment and increased overall yields compared to 2016. The use of improved agronomic package of ridging, yam seed pre-treatment, trellis staking and fertilizer rate of 30:30:36 N: P_2_0_5_:K_2_0 kg/ha plus 15 kg/ha Mg and 20 kg/ha S as MgSO_4_ resulted in total tuber yield percentage difference over farmers’ practice of mounding and planting of 68.6%, 78.3%, 80.7% for the 2015 cropping season and 113.6%, 113.9%, 120% for 2016 cropping season in Ejura, Atebubu and Kintampo farming communities (Fig 5) respectively. Stand count/ha on the ridges were significantly (P<0.01) higher on the improved agronomic practice fields than farmers’ practice fields for all the locations and across season (Figs 2–4). This we attribute to the use of ridges which made it possible to plant at 1.2m between ridges and 0.8m on the ridges resulting in a planting density of about 10,416 stands per hectare whiles the farmers practice of mounding were relatively sparse (1.5m-2m) resulting in just about 3,400 - 4,000 stands/ha. Thus, ridging helped maintain optimum number of plants and promoted efficient use of fertilizer and conserved moisture than mounding.

**Figure 2:**
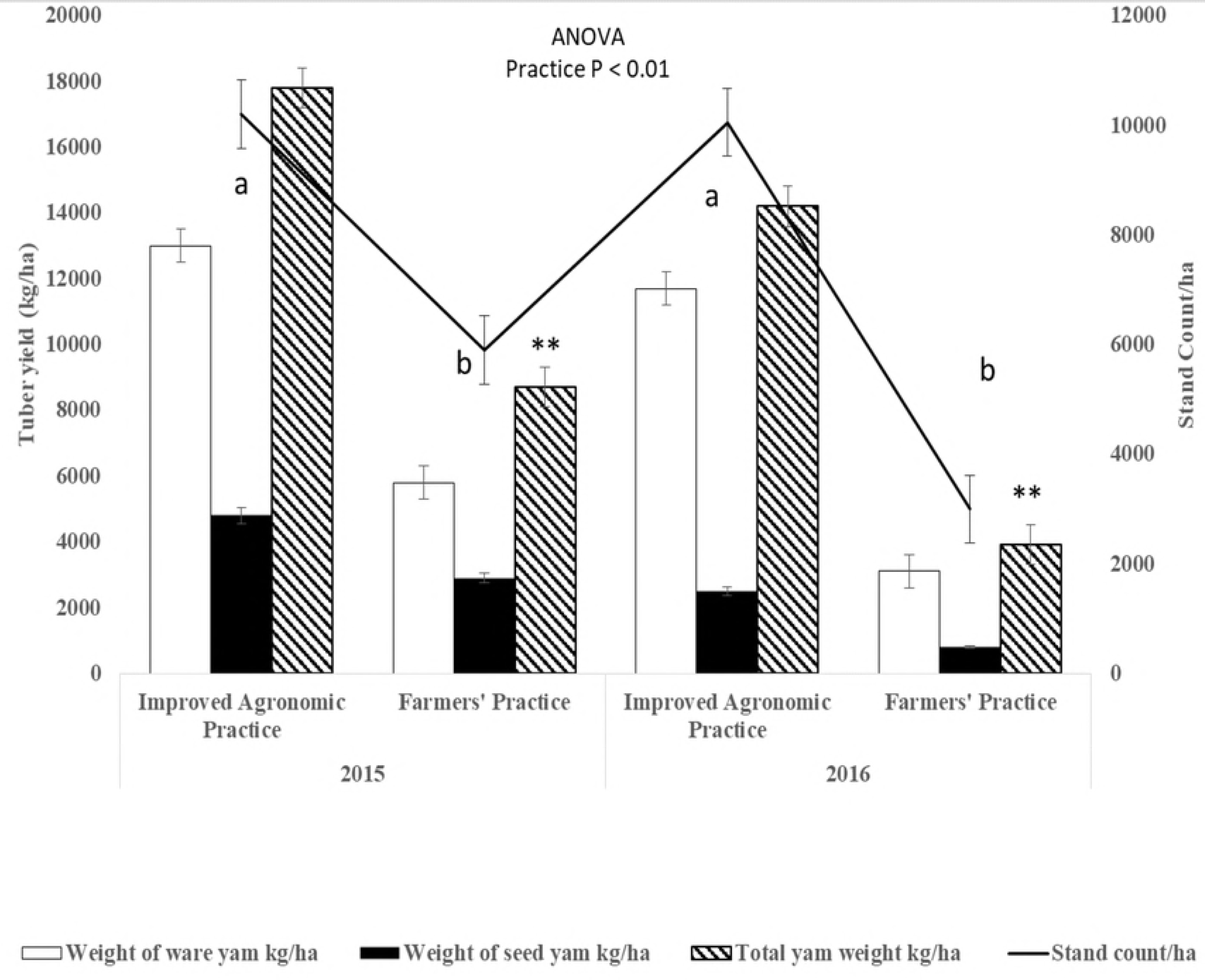
Yam tuber yields and stand count as influenced by improved agronomic practice and farmers’ practice in the Ejura farming communities for 2015 and 2016 cropping seasons. Mean values and standard errors (n = 4) are plotted. Index letters above the bars indicate significant differences (P < 0.01) between media not sharing the same letter by Tukey’s HSD test. Asterisks indicate significant differences (P < 0.01) between the two practices.

**Figure 3:**
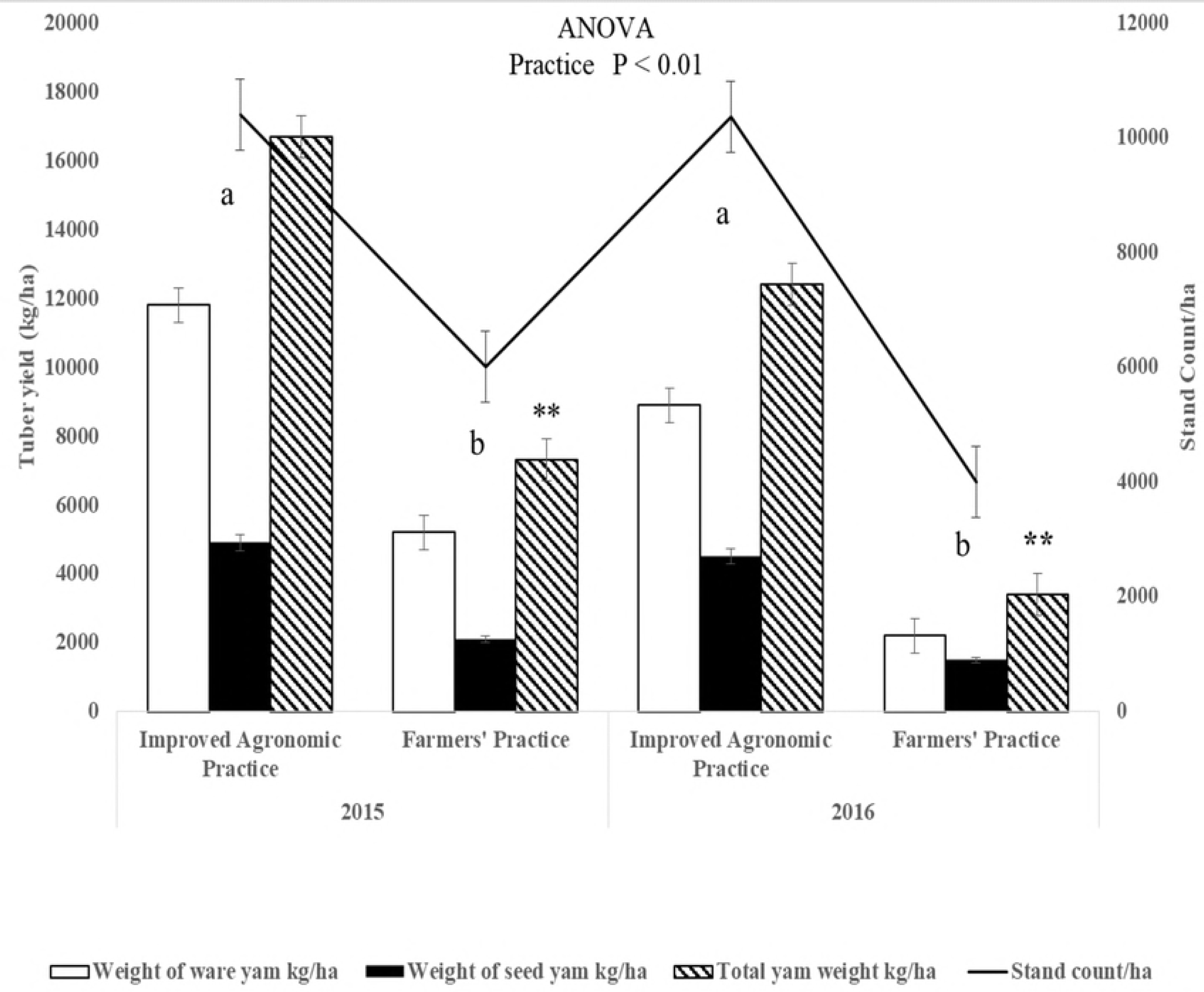
Yam tuber yields and stand count as influenced by improved agronomic practice and farmers’ practice in the Atebubu farming communities for 2015 and 2016 cropping seasons. Mean values and standard errors (n = 4) are plotted. Index letters above the bars indicate significant differences (P <0 .01) between media not sharing the same letter by Tukey’s HSD test. Asterisks indicate significant differences (P < 0.01) between the two practices.

**Figure 4:**
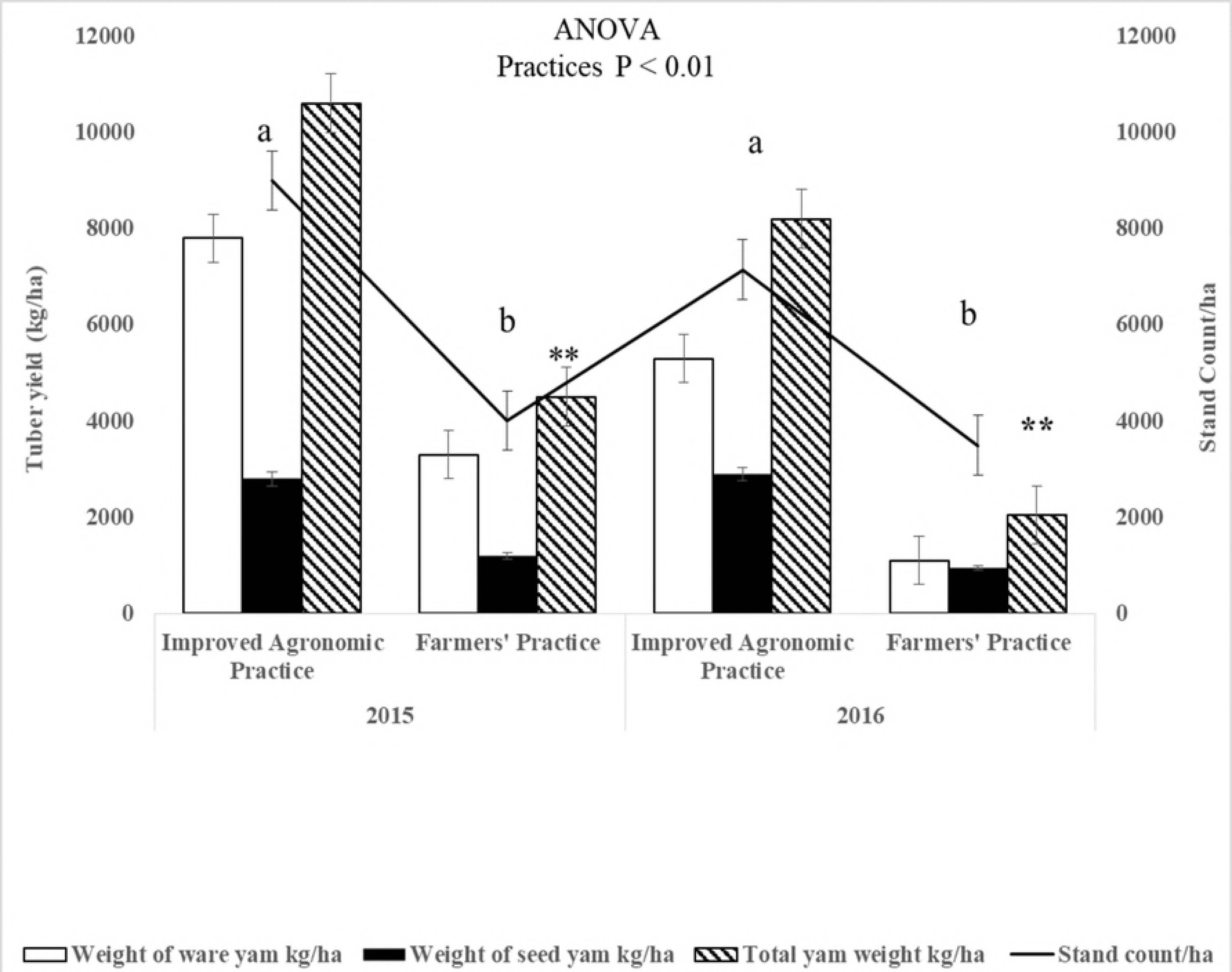
Yam tuber yields and stand count as influenced by improved agronomic practice and farmers’ practice in the Kintampo farming communities for 2015 and 2016 cropping seasons. Mean values and standard errors (n = 4) are plotted. Index letters above the bars indicate significant differences (P < 0.01) between media not sharing the same letter by Tukey’s HSD test. Asterisks indicate significant differences (P < 0.01) between the two practices.

**Figure 5:**
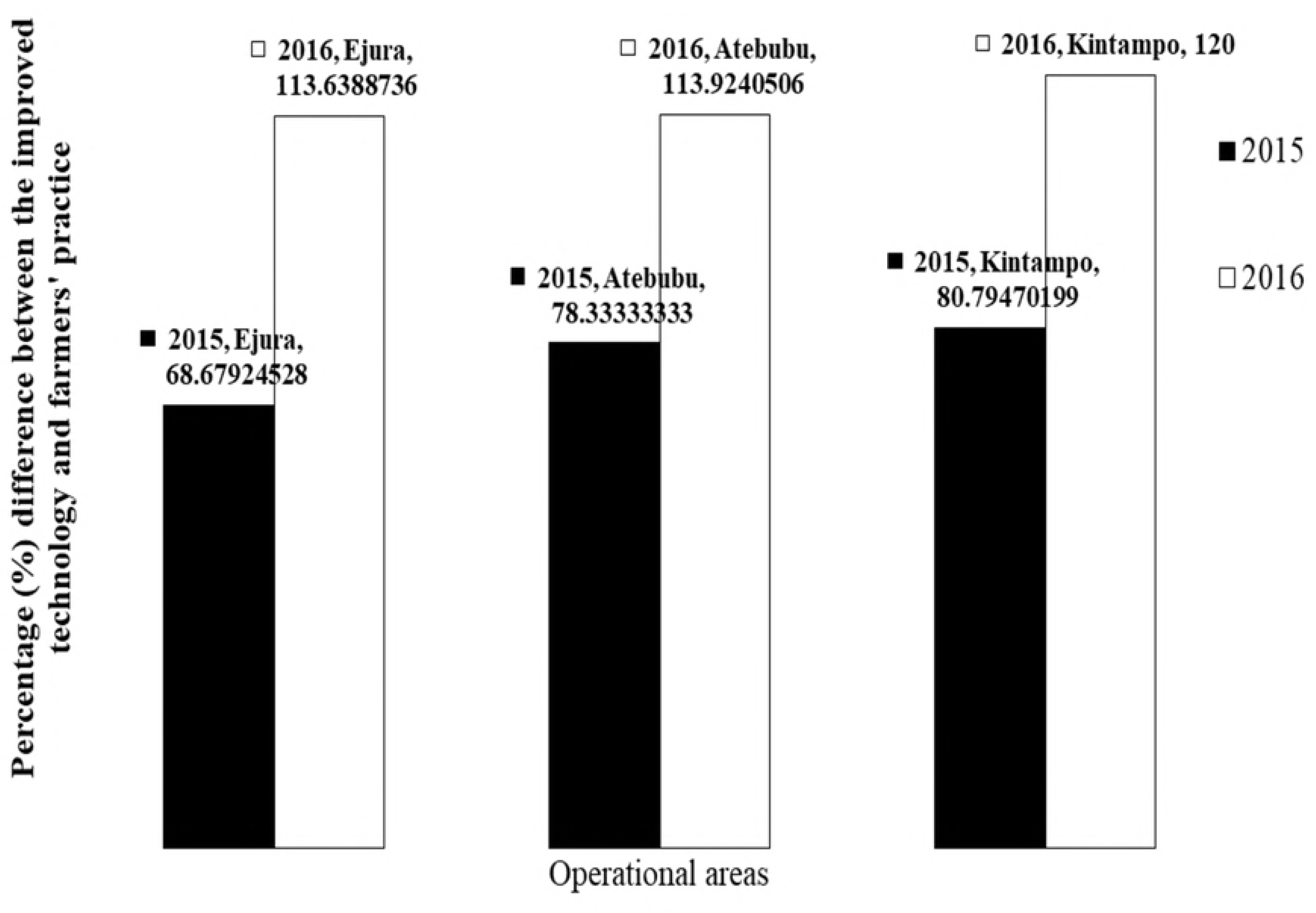
Percentage differences between the two practices across two seasons calculated for each location based on their total yam yield (kg/ha).

It was observed that the improved agronomic fields had better yam canopy as a result of the combined effect of trellis staking and ridging. Revealing similar tuber yield trends on fertilized mounds and unfertilized ridges (8) supports the argument that fertilizer application helps the farmer to achieve value by being more efficient and profitable on ridges than on mounds. Moreover, pre-treating seed yam before planting on the improved agronomic fields resulted in a reduction in yam rot and increased stands/ha than on the farmers’ practice fields where seeds were not treated resulting in lower stands/ha. Seed treatment for pest and disease before planting is recommended to promote sprout rate ensuring improvement in final stand density culminating into overall high yam yields (11,12,17). Erratic rainfall and prolong drought require technologies that enable the soil to conserve moisture and promote nutrient use efficiency in order to increase resiliency of any cropping system. An observation we made during the studies was that drought was more pronounced on the farmers’ practice which used relatively sparse mounds and vertical staking option than on the improved agronomic practices which used ridges and trellis staking option. Similar studies suggest that planting on ridges can maintain optimum plant stands and conserve more moisture than mounds resulting in more efficient water used on ridges than on mounds (7,8,17,21). These attributes makes the use of ridges more soil nutrient efficient upon application of fertilizer than on mounds. Furthermore, (8) made a similar observation with planting on ridges increasing yields significantly than on mounds.

It is recommended that adopting and following through the improved agronomic package based on results of 2016 cropping season (Figs 1–5) where rainfall were considerably lower illustrates the resilience ability of it during reduced precipitation. In spite of the gains in yam fertilization, there are perceptions and claims by some consumers and farmers in the public space that fertilizing yam leads to rots and reduces the overall shelf life. We recommend further research into these claims as we settled the dust in this paper (Table 3) on claims that fertilizer affected the taste quality of yam.

### Partial budgeting and cost benefit analysis of the two practices

**Table 2** presents the partial budgeting and cost benefit analysis of "*Dente"* white yam production under improved agronomic practice and farmers’ practice in Ejura, Atebubu and Kintampo operational areas. The results revealed that irrespective of location or the season, yam planted with the improved agronomic package had higher benefit to cost ratio compared to farmers’ practice. Benefit to cost ratios of 4.76:3.64, 4.04:2.48 and 2.01:1.08 were achieved for Ejura, Atebubu and Kintampo communities respectively for the 2015 cropping season (**Table 2**) in sequence of improved technology: farmers’ practice. Thus, when a farmer invests Gh₵ 1.00 in yam production using the recommended improved technology a profit of Gh₵ 3.76, Gh₵ 3.04 and Gh₵ 1.01 was to be accrued in addition to the Gh₵ 1.00 invested capital at Ejura, Atebubu and Kintampo respectively during the 2015 season. During the 2016 cropping season, drought was more intense particularly for Atebubu and Kintampo areas (Figure 1). This however did not affect benefit to cost ratio for using the improved agronomic package thus achieving 3.76, 3.03 and 1.33 compared to 1.16, 0.75 and −0.55 for Ejura, Atebubu and Kintampo communities respectively (**Table 2**). Thus, a profit of Gh₵ 2.76, Gh₵ 2.03 and Gh₵ 0.33 was to be accrued in addition to the Gh₵ 1.00 invested capital upon the use of improved agronomic practices at Ejura, Atebubu and Kintampo respectively. The use of the farmers’ practice resulted in total loss of Gh₵ 1.55 in Kintampo area (**Table 2**). This suggest that the use of the improved agronomic practice would not only increase and sustain yields on continuously cropped fields but also the it is the best option during drought spells, erratic and reduced rainfall conditions. The improved agronomic package thereby increases farmer’s resilience in dealing with such harsh weather conditions with assured returns on their investments.

### Influence of fertilizer on the taste of boiled yam

Dente yam planted in 2016 under the improved agronomic practice (fertilized) and farmers’ practice (unfertilized) were boiled after harvest and given to fifty participants each from Atebubu, Ejura and Kintampo who were mainly farmers for sensory evaluation (Table 3). Participants were not previewed as to whether the yam they evaluated at a given time was fertilized or unfertilized as they were coded in order to avoid bias. Participants assessed the various boiled yam on three culinary qualities: ‘taste’, ‘texture’ and ‘aroma’ (Table 3) based on their individual preferences from a scale of 1 up to 5 with 1 been the best score and 5 as the worst. Overall acceptability and STD acceptability on the three traits; taste, texture and aroma was subsequently calculated following the approach of (8). The results was in line with previous results by (8) that, contrary to the view that the use of fertilizer in yam production affects the quality of yam, sensory evaluation showed that the culinary qualities of fertilized yam is good and could even be better than unfertilized yam.

## Conclusion and Policy Implication

The improved agronomic technology has proven to be climate resilient comparing the overall yield (significantly >60% difference) trends of 2015 and 2016 seasons against farmers’ practice given the rainfall amounts and pattern in those years. The overarching difference between what farmers do today and the improved agronomic model is the intensification drive and a higher use efficiency (staking, nutrient, soil moisture conservation, improved sprouting) of the technology at a given area as a consequence of the high planting density allowed by the linear arrangement of ridging, trellis staking and seed treatment. It is anticipated that more and more farmers who have shown keen interest would adopt the technology through the gains made. Claims on yam fertilization affecting taste quality proved otherwise with even a higher overall acceptability. Further research is needed to understand claims and perceptions by farmers on fertilizer affecting yam storage. Up-scaling of the improved agronomic technology to other yam growing communities would further augment the gains already made.

## Acknowledgement

We thank the Bill and Melinda Gates Foundation who funded this research work through YIIFSWA project that was coordinated by IITA, Nigeria through CSIR-Crops Research Institute. We are grateful to Mr. Henry Azotibah (Ejura), Mr. Kyeremeh (Atebubu) all of MoFA and Mr. Enoch Bessah of CSIR for their invaluable support and advice during data acquisition and analysis.

## Abbreviations

CSIR: Council for Scientific and Industrial Research
FAO: Food and Agriculture Organisation
IITA: International Institute for Tropical Agriculture
MoFA: Ministry of Food and Agriculture
YIIFSWA: Yam Improvement for Income and Food Security in West Africa

